# Multipotent ubiquitin/ubiquitin-like deconjugation activity of the *rhizobial* effector NopD

**DOI:** 10.1101/2023.09.20.558580

**Authors:** Ying Li, Jordi Perez-Gil, Maria Lois, Nathalia Varejão, David Reverter

## Abstract

Post-translational modification of proteins by ubiquitin-like modifiers (UbL), such as SUMO, ubiquitin or Nedd8, contribute to regulate most pathways in the cell. Protein modification can be reversed by dedicated UbL deconjugating proteases families. During bacterial infection, a repertoire of effector proteins, including deconjugating proteases, are released to perturb the host cell defense to favor bacterial survival. NopD, an effector protein from rhizobia involved in nodule symbiosis in legumes, possesses deSUMOylation, and unexpectedly, deubiquitination and deNeddylation activities. Here we show two crystal structures of *Bradyrhizobium* NopD in complex with either *Arabidopsis* SUMO2 and ubiquitin at 1.50 or 1.94 Å, respectively. Despite their low sequence identity, SUMO and ubiquitin interact with a similar NopD interface by means of a unique loop insertion in the NopD sequence. Biochemical and infiltrations in tobacco leaves reveal specific residues that discriminate between deubiquitination and deSUMOylation. These unusual multiple deconjugating activities against SUMO, ubiquitin and Nedd8, represent a paradigmatic example of an optimized protease to perturb distinct UbL post-translational modifications during host cell infection.

**Significance Statement:** During bacterial infection, including *rhizobia* symbiosis in legume plants to fix atmospheric nitrogen, a set of effector proteins, such as ubiquitin/ubiquitin-like deconjugating proteases, are released to perturb the host cell defense to favor bacterial survival. We have discovered that the *rhizobial* effector protein, NopD, encompasses triple deconjugation activities against SUMO, ubiquitin and Nedd8. Structural analysis of NopD in complex with SUMO and ubiquitin reveals the presence of a loop insertion in the protease-substrate interface to allow this multiple substrate binding capability. Such unusual deconjugating activities in NopD for ubiquitin, SUMO and Nedd8 modifiers, represent a paradigmatic example of an optimized protease domain to perturb distinct UbL post-translational modifications during host cell infection.

## Introduction

Bacterial infection and colonization in eukaryotic host cells requires special mechanisms to shut down host defense systems and ensure pathogen or symbiont survival. A common strategy by infectious bacteria consist on the injection of a plethora of effector proteins into the eukaryotic cell host by complex multiprotein structures, such as the needle-like Type III secretion system (T3SS), which spans the inner and outer bacterial membranes and is highly conserved among bacterial pathogens ^1^. T3 protein effectors are injected by pathogenic bacteria into eukaryotic host cells to hijack the host defense systems. In plants, in addition to pathogenicity, T3SS has also been identified in rhizobia, a symbiotic bacteria that resides in legumes and can fix the atmospheric nitrogen to ammonia by the formation of nodules in the roots of legume plants ^2,3^.The effector proteins of the rhizobia T3SS are called Nops (Nodulation Outer Protein), and around 20 to 30 T3 effector proteins can be secreted into the legumes host cells, such as NopL ^4^, NopE1/E2 ^5^, NopM ^6^, NopT ^7^ and NopP ^8^. Signaling kinase cascades or the eukaryotic UPS (Ubiquitin-Proteasome system) are pathways often targeted in the host by T3 bacterial effectors to allow infection ^9^.

Ubiquitin can regulate a large number of cellular processes in the eukaryotic cells by the formation of distinct polymeric chains, including protein half-life, localization, endocytosis, autophagy, and many more major functions ^10^. Ubiquitin is ubiquitous and highly conserved in eukaryotes, including animals, yeast and plants. Additionally, Ubiquitin-like modifiers (Ubl), such as SUMO, Nedd8 or ISG15, can also regulate many cellular pathways by covalent modification of protein targets ^11^. In general, UbL modification (ubiquitin, SUMO or Nedd8) is conducted by an enzymatic cascade orchestrated by an E1 activating enzyme, an E2 conjugating enzyme, and E3 ligases ^12^. Ubiquitin or SUMO modification can be reversed by the action of several dedicated families of proteases called deubiquitinates (DUBs) and deSUMOylases (SENP/ULP). Humans contain around 100 DUBs grouped in 6 different families based on their structure and action mechanisms ^13^. Most DUBs are specific for ubiquitin, except a few exceptions that can cleave off ISG15 (USP18) or SUMO (USPL1) from protein targets ^14^. The deSUMOylase CE clan of cysteine proteases in humans consists of six SUMO-specific proteases (SENP1-3 & 5-7) and one NEDD8-specific protease (NEDP1/SENP8). So far, all members of the eukaryotic DUBs or SENP/ULP families are specific for deconjugation of only one type of UbL ^15^.

Also, a number CE family enzymes are encoded in pathogenic bacteria and in bacteria that reside inside eukaryotic cells (plant symbiont) ^2,3^. Since no Ubl-type modifier systems exist in these bacteria, they are injected as effectors to manipulate the eukaryotic host signaling pathways. Some of these CE family effectors include the ChlaDUB1 in *Chlamydia* ^16^, LegCE of *Legionella* ^17^, SseL of *Salmonella* ^18^, XopD of *Xanthomonas* ^19^, and ElaD of *E. coli* ^20^, RickCE from *Ricksettia* ^17^ or ShiCE from *Shigella* ^17^. Interestingly, most bacterial CE proteases are specific for ubiquitin instead of SUMO, despite their structural similarity with the SENP/ULP family ^17^. In some cases they even possess an unusual Ser/Thr acetyltransferase activity, such as YopJ from *Yersinia* ^21^. Structural comparison between bacterial protease effectors from different organisms indicate a high versatility of the CE catalytic domain to acquire particular functions ^17^, such as the capability to cleavage K63 or K48 polyubiquitin chain linkages.

Recently in rhizobia a novel T3 effector was identified belonging to the CE protease clan with specific activity against plant SUMOs ^22^. NopD, from *Bradyrhizobium sp.* XS1115, contains a C-terminal CE protease domain with sequence similarity to XopD, a T3 effector from the plant pathogen *Xanthomonas campestris* ^23^. Interestingly, *Xanthomonas* XopD has the unique property to possess deSUMOylating and deubiquitinating activities by the formation of two different interfaces to cleave either SUMO or ubiquitin ^17^. This dual activity for SUMO and ubiquitin is quite unusual feature in eukaryotic DUBs and inSENP/ULP family members.

A recent characterization of rhizobia NopD indicate that the CE protease activity is required for cell death induction in tobacco and it can reduce the size of the nodule formation in a model legume plant *Tephrosia vogelii* ^22^. Also, several NopD-like effector proteins have been identified in different rhizobia species possessing a CE catalytic protease domain similar to XopD ^24^, such as Bel2-5 in *Bradyrhizobium elkanii*, which seems to have a role in root nodule formation ^25^, or MA20_12780 from *Bradyrhizobium japonicum*, also suggested to act as a virulence factor in nodulation ^26^. In all instances, the deconjugation activity of T3 NopD-like effectors have a major role during rhizobia infection and formation of nodules in legume roots.

In this work, we reveal that the T3 effector protein of *Bradyrhizobium*, NopD, in addition to deSUMOylating activity, also encompasses K48 linkage-specific deubiquitinating and deNeddylating activities, which it is quite unusual since these proteases are generally specific for a single type of UbL modifier. We have determined the molecular mechanism for such uncommon deconjugation activities by solving the crystal structures NopD in complex with either *Arabidopsis* SUMO2 or ubiquitin at 1.5 or 1.9 Å resolution, respectively. Exceptionally, in both complexes NopD utilizes a similar binding interface despite the low sequence identity between SUMO and ubiquitin. Biochemical assays and in vivo tobacco leaves infiltrations have revealed the particular residues in the NopD interface that are responsible for the deSUMOylating and deubiquitinating activities.

## Results

### NopD has an unusual multiple activity for SUMO, ubiquitin and Nedd8

Structural predictions indicate that the 1017 residue full-length protein of the *Bradyrhizobium* NopD symbiotic effector consists of a long N-terminal extension, which seems disordered based on Alphafold-2 models, followed by a globular C-terminal protease domain with homology with the CE clan of cysteine proteases. This C-terminal catalytic domain was reported to possess deconjugating activity for plant SUMOs ^22^, displaying the highest homology with XopD, a bacterial plant effector protease from *Xanthomonas campestris* (22.3% sequence identity for 184 residues) ^23^. The catalytic domain of NopD also displays low homology with the eukaryotic ULP/SENP deSUMOylase family. So, based on sequence alignments with members of the SENP/ULP family (Supplementary Figure 1), we generated a C-terminal domain construct of NopD (from Gly832 to Asn1016) that was expressed in *E. coli*.

*Xanthomonas* XopD was reported to possess a dual deconjugation activity for SUMO and ubiquitin ^17^, a quite unusual property in the SENP/ULP family. Surprisingly, we observe that *rhizobium* NopD also displays deconjugation activity for both plant SUMO and ubiquitin (Figure 1a). NopD binds *Arabidopsis* SUMO2-PA or ubiquitin-PA probes, in contrast to the lack of binding to human SUMO2-PA. The specificity of the interaction was demonstrated by the absence of crosslinked adducts when catalytic inactive active site cysteine to alanine mutants (C971A) was checked (Figure 1a).

**Figure 1.**
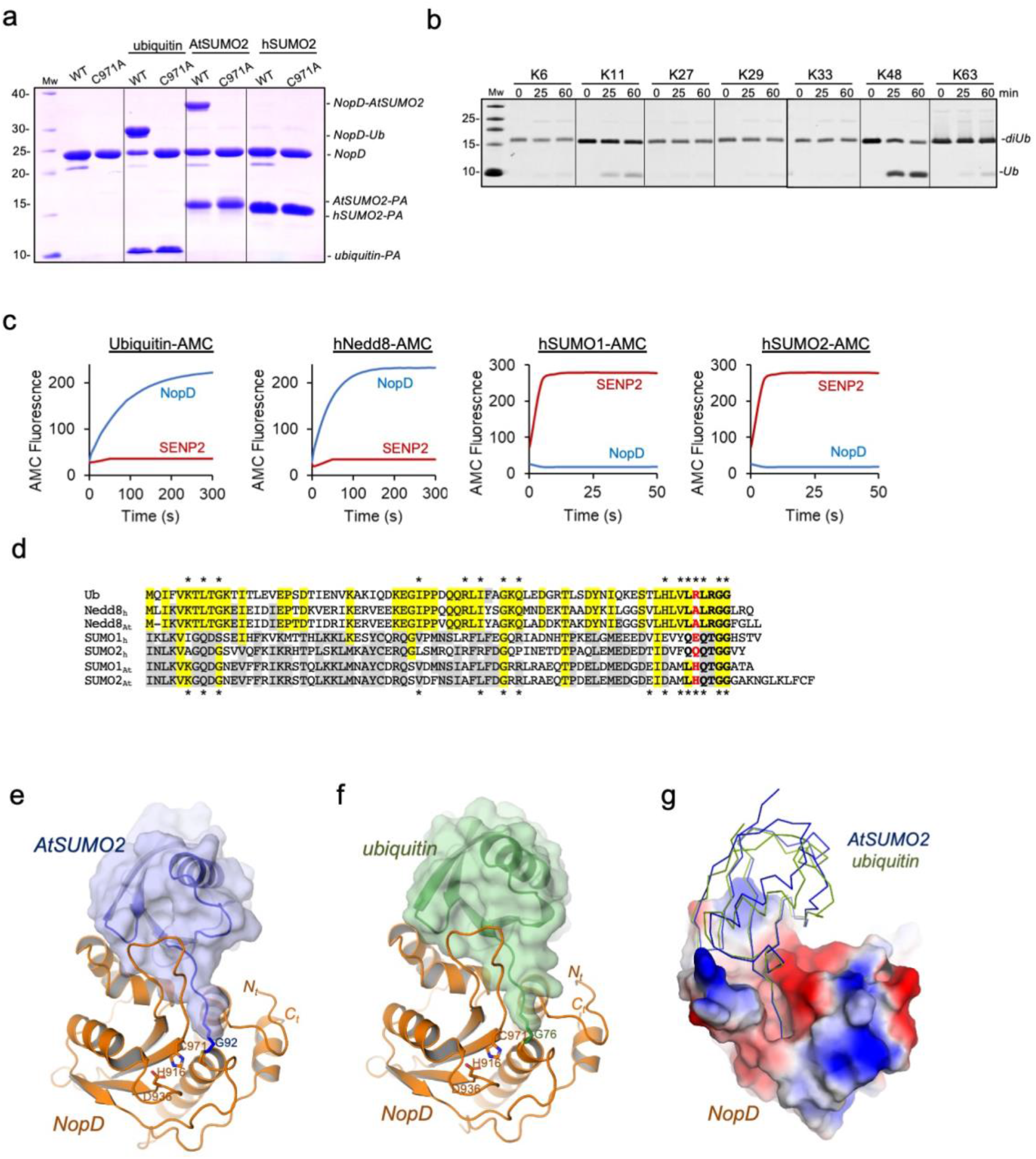
Specificity of NopD for *Arabidopsis* SUMO, ubiquitin and Nedd8. **a,** SDS-PAGE of the binding between the NopD wild-type and active site mutant C971A against ubiquitin-PA, *Arabidopsis* SUMO2-PA and human SUMO2-PA suicide probes for 2 hours**. b,** SDS-PAGE of the di-ubiquitin linkage specificity analysis for NopD over time. The concentration of NopD and diUb are 600nM and 3 μM, respectively**. c,** Time-course plots of fluorescent AMC-based substrates (100 nM) of ubiquitin, human Nedd8, human SUMO1 and SUMO2 substrates with the catalytic domains of *rhizobia* NopD (25 nM) and human SENP2 (25 nM) as a control. Reactions were conducted in triplicate and the average curve is displayed. **d,** Sequence alignment of ubiquitin, human and *Arabidopsis* Nedd8 and, human and *Arabidopsis* SUMO1 and SUMO2. Yellow and grey background depict sequence conservation between ubiquitin and Nedd8, and between SUMO isoforms, respectively. The C-terminal tails before the diGly motif are marked in black bold letters. Ubiquitin Arg72 position in marked as red bold letter. Asterisks indicate binding residues of NopD with either *Arabidopsis* SUMO2 (blue) or ubiquitin (green), respectively. **e,** Cartoon representation of the NopD catalytic domain with AtSUMO2. **f,** Cartoon representation of the NopD catalytic domain with ubiquitin. Catalytic triad residues are label and shown in stick representation. The N- and C-terminal are labeled. **g,** Electrostatic potential surface representation of NopD in complex with overlapped ribbon structures of *Arabidopsis* SUMO2 (blue) and ubiquitin (green).

Next, the NopD preference for the different types of poly-ubiquitin chains was assessed by using a di-ubiquitin chain assay kit (Figure 1b). Whilst the CE clan of bacterial effectors with deubiquitinating activity, some present in human, normally display specificity for K63-linked chains, in plants, the unique known pathogen *Xanthomonas* XopD displayed preference for K48, and to a lesser extend to K11 and K29 chains ^17^. Similarly, in symbiotic *rhizobia,* NopD also displays a preference for K48 ubiquitin linkages, and to a lesser extent for K11 linkages (Figure 1b).

Interestingly, in addition to plant SUMO and ubiquitin, we show that *rhizobia* NopD also exhibits a strong deconjugating activity for a different type of UbL modifier, such as Nedd8 (Figure 1c). NopD displays strong activity against Nedd8 and ubiquitin using fluorescent-AMC substrates, but it is inactive for human SUMO1 or SUMO2. In contrast to the human deSUMOylase SENP2, which is only active for human SUMO1 and SUMO2 (Figure 1c). The proteolytic activity for Nedd8 is unique to NopD and was not observed in *Xanthomonas* XopD ^17^. Sequence alignment shows lower identity between human or plant SUMOs with either ubiquitin or Nedd8, being the latter two quite similar (Figure 1d). The presence of a deconjugating protease, such as *rhizobia* NopD, which is simultaneously active for ubiquitin, Nedd8 and SUMO is quite uncommon, and might represent a paradigmatic example of an optimized deconjugating protease with multiple activities during host infection.

### Overall complex structures of NopD with *Arabidopsis* SUMO2 and ubiquitin

To form complexes of NopD with either *Arabidopsis* SUMO2 and ubiquitin, the C-terminal carboxylate groups of SUMO and ubiquitin were chemically modified with a highly reactive and stable C-terminal alkyne (SUMO2-PA or ubiquitin-PA) after a chemical reaction with propargylamine ^27–29^. The C-terminal carboxylate group of SUMO2-PA or ubiquitin-PA can thus be crosslinked with the active site cysteine of NopD (Cys971) to form a stable covalent product complex after incubating NopD with SUMO2-PA or ubiquitin-PA. In an initial screening, a few diffraction quality crystals of NopD-SUMO2 or NopD-ubiquitin were produced in conditions containing 0.1 M imidazole (pH 7.0), 50% MPD or 0.1 M imidazole (pH 8.0), 10% PEG8000, respectively. Molecular replacement with human ubiquitin (PDB 1UBQ), *Arabidopsis* SUMO2 and NopD Alphafold-2 models assisted to determine the complex structures. Two complexes per asymmetric unit were found in the NopD-SUMO2 crystals, which belonged to the P2_1_2_1_2_1_ space group and diffracted to a 1.50 Å resolution; and one complex per asymmetric unit in the NopD-ubiquitin crystals, which belonged to the P4_3_ space group and diffracted beyond 1.94 Å resolution (Table 1).

**Table 1.** Crystallographic statistics of NopD-AtSUMO2 and NopD-Ubiquitin crystal structures.

|  | *NopD-AtSUMO2* | *NopD-ubiquitin* |
| --- | --- | --- |
| **Data collection** |  |  |
| Space group | P2<sub>1</sub>2<sub>1</sub>2<sub>1</sub> | P4<sub>3</sub> |
| Unit cell parameters (Å) | 80.85, 86.83, 90.49 | 86.50, 86.50, 46.67 |
| Wavelength (nm) | 0.9791 | 0.9792 |
| Resolution range (Å) | 49.52 – 1.50 | 43.25 – 1.94 |
| R<sub>merge</sub> | 0.07 (0.83) | 0.07 (1.42) |
| R<sub>pim</sub> | 0.03 (0.48) | 0.03 (0.60) |
| (I/σ(I)) | 14.2 (1.8) | 13.3 (1.2) |
| Completeness (%) | 92.7 (48.9) | 94.2 (55.0) |
| Multiplicity | 6.1 (3.7) | 6.8 (6.6) |
| CC (1/2) | 0.99 (0.49) | 0.99 (0.51) |
| **Structure refinement** |  |  |
| Resolution range (Å) | 49.52-1.50 | 43.25-1.94 |
| No. of unique reflections | 76875 | 22770 |
| R<sub>work</sub> / R<sub>free</sub> (%) | 17.84 / 20.36 | 17.16 / 19.94 |
| No. of atoms |  |  |
| Protein | 8363 | 2055 |
| Water molecules | 312 | 62 |
| Overall B factors (Å<sup>2</sup>) | 31.02 | 59.74 |
| Rms deviations |  |  |
| Bonds (Å) | 0.015 | 0.007 |
| Angles (°) | 1.140 | 0.839 |
| Ramachandran favored (%) | 99.22 | 98.41 |
| Ramachandran allowed (%) | 0.59 | 1.19 |
| Ramachandran outliers (%) | 0.2 | 0.4 |

The final models of NopD displayed the canonical fold of CE cysteine protease clan and contained a continuous sequence from Pro829 to Ala1011 in complex with SUMO2, and from Gly828 to Leu1009 in complex with ubiquitin (Figure 1e & 1f). The final electron density maps clearly show the covalent bond formed between the SUMO or ubiquitin C-terminal glycine and the NopD active site Cys971, confirming the specificity of the catalytic reaction between NopD with SUMO2-PA or ubiquitin-PA substrates (Supplementary Figure 2). Unexpectedly, both SUMO and ubiquitin interact with a similar surface of NopD (Figure 1g), in contrast to the different interfaces observed in complex with *Xanthomonas* XopD ^17^.

Structural overlapping with the unbound NopD AlphaFold-2 model is quite similar, displaying mainchain rmsd values (root mean square deviation) of 0.93 Å and 0.90 Å, respectively (Supplementary Figure 3). Only residues involved in the interface with SUMO/ubiquitin display different orientations, such as the “Loop insert” located between β2 and β3 strands (Figure 2a and Supplementary Figure 3). The active site catalytic triad seems already formed in the absence of SUMO or ubiquitin substrate, as deduced by comparing the distances between the catalytic triad residues (Cys971, His916 and Asp936) with the AlphaFold-2 model, suggesting that NopD might be active in the absence of a UbL substrate, as initially observed in human SENP2 ^30^, and does not need any substrate-induced activation mechanism.

**Figure 2.**
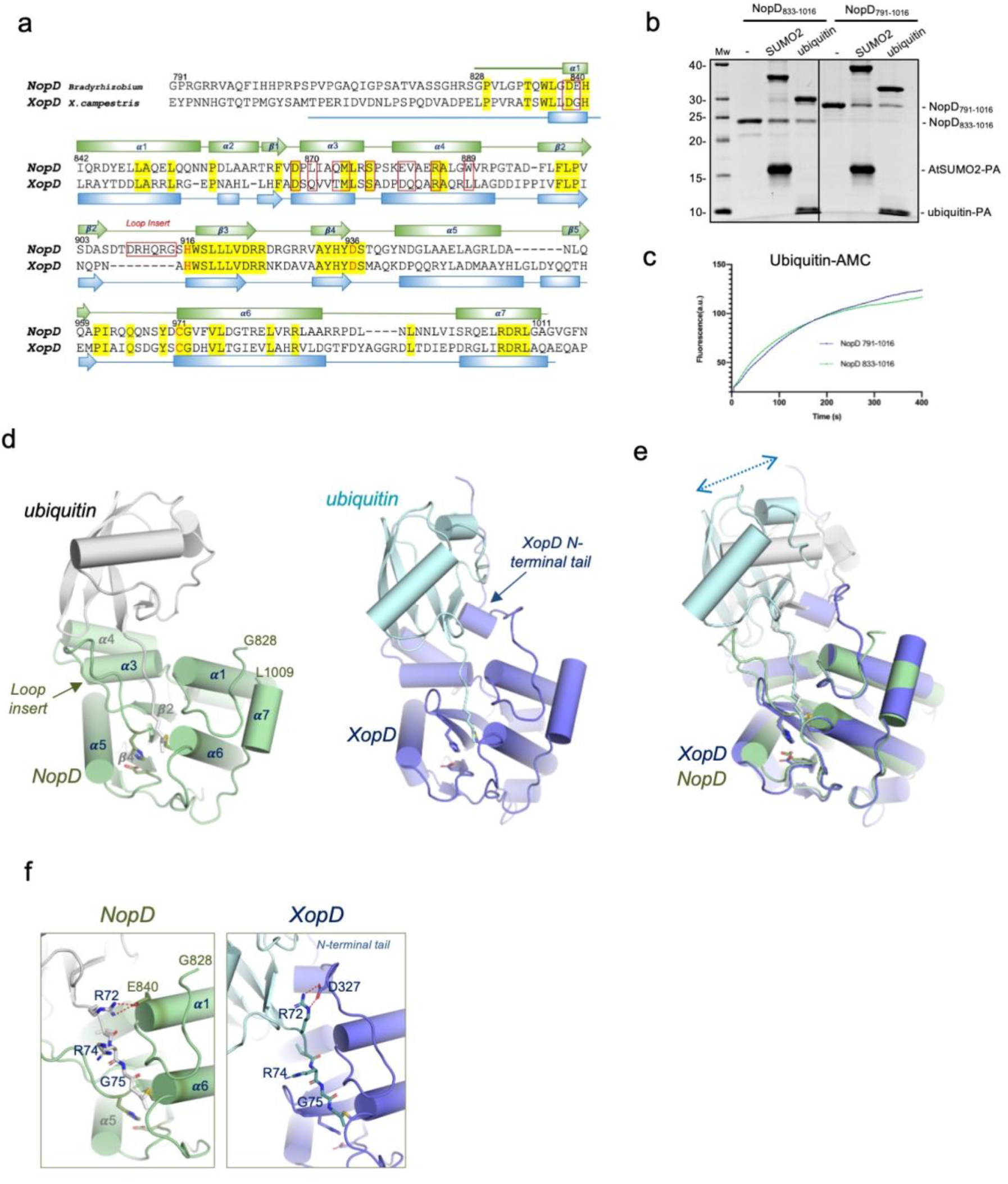
The deubiquitinating activity of NopD does not depend on an N-terminal extension. **a,** Structural alignment of sequences corresponding to the catalytic domains for NopD and *Xanthomonas campestris* XopD. Red squares indicate interface residues to NopD. Catalytic triad residues are shown in red. Secondary structure cartoon is depicted above for NopD (green), or below for XopD (blue). **b**, SDS-PAGE of binding of NopD with a longer N-terminal extension against AtSUMO2-PA and ubiquitin-PA suicide probes for 2 hours. Bands of NopD crosslink, NopD_791-1016_, NopD_833-1016_, AtSUMO2-PA, ubiquitin-PA are labeled. **c,** Time-course of ubiquitin-AMC hydrolysis for NopD_833-1016_ and NopD_791-1016_ constructs. Ub-AMC (100 nM) was incubated with NopD (5 nM) constructs over time. Reactions were conducted in triplicate and the average curve is displayed. **d,** Cartoon representation of the NopD-Ubiquitin (green) and *Xanthomonas campestris* XopD-Ubiquitin (PDB: 5JP3) complexes. The catalytic residues are shown in stick representation. Loop insert in NopD is labeled and marked. XopD N-terminal tail extension is indicated**. e**, Superposition of the *Xanthomonas* XopD-Ubiquitin (PDB: 5JP3) complex (blue) with NopD-ubiquitin complex (green). Double headed discontinued blue arrow indicates the displacement of ubiquitin between NopD (grey) and XopD (light blue) complexes. **f**, Close-up views of the C-terminal tails of ubiquitin in NopD-ubiquitin complex (green) and in the XopD-ubiquitin complex (purple). Relevant residues are labeled and shown in stick representation. Ubiquitin Arg72 electrostatic interactions are represented by dashed red lines.

### Binding interface of NopD with ubiquitin and SUMO

The *Xanthomonas* XopD-ubiquitin structure (PDB 5JP3) revealed a novel interface between a N-terminal extension of XopD with the Ile44-patch of ubiquitin ^17^. This N-terminal extension in XopD was essential for the deubiquitinase activity, and full deletion or single point mutations caused failed binding to ubiquitin-PA. Accordingly to the *Xanthomonas* XopD sequence, we produced a NopD catalytic domain with a longer N-terminal extension to check its relevance in deubiquitinating assays. Two NopD constructs were produced: a longer domain from Gly791 to Asn1016, and a shorter domain from Pro833 to Asn1016 (Figure 2a).

In contrast to *Xanthomonas* XopD, the N-terminal extension in NopD does not play a significant role in the catalytic activity against ubiquitin substrates, displaying similar binding properties and catalytic activities in both NopD constructs (Figure 2b & 2c). Consequently, a different interface takes place in the NopD complex with ubiquitin, as observed in the structural overlap of NopD/XopD complexes with ubiquitin (Figure 2d & 2e), displaying a notable displacement of ubiquitin around 5-6 Å. Interestingly, the ubiquitin interface in NopD is similar to the interface with AtSUMO2, despite the lower identity between SUMO and ubiquitin (Figure 1g and Supplementary Figure 4).

An interesting feature in the NopD-ubiquitin complex is the presence of a strong well-oriented electrostatic interaction between ubiquitin’s C-terminal Arg72 and NopD Glu840 (Figure 2f). This interaction is unique in NopD, since the equivalent position in XopD contains a glycine (Figure 2a), and in XopD ubiquitin’s Arg72 interacts with the N-terminal extension residue Asp327 (Figure 2f). Glu840 probably might be a particular acquisition of NopD to interact with ubiquitin in symbiotic *Rhizobium (*Supplementary Figure 5). Ubiquitin’s C-terminal tail contains two arginine residues (Figure 1d), whilst Arg74 is not engaged in any specific contact, Arg72 is usually involved in specific electrostatic contacts with deubiquitinating enzymes ^31^. Later in Figure 5, our in vitro assays confirm the major role of the Arg72-Glu840 interaction in the ubiquitin complex.

Moreover, another unique characteristic in the NopD structure is the presence of a “Loop insert” between strands β2 and β3 (Figure 2a & 2d). The only other member in the eukaryotic SENP/ULP family with a similar insertion corresponds to the SENP8/NEDP1 member, which cleaves off Nedd8 instead of SUMO ^32,33^. Intriguingly, this “Loop insert” participates in the interaction with the C-terminal tail of both ubiquitin and SUMO, and it is essential for binding and proteolytic activity of NopD with SUMO and ubiquitin (later in Figure 5).

### C-terminal tail comparison of SUMO2 and ubiquitin in NopD

The C-terminal tails of ubiquitin and SUMO2 are buried in a NopD surface cleft that contains the active site catalytic triad, responsible for cleaving off the isopeptidic bond after the diGly motif in both SUMO and ubiquitin C-terminus. Several conserved contacts are observed despite the low-identity of the C-terminal tails of AtSUMO2 (-MLHQTGG) and ubiquitin (-VLRLRGG). Similar to other members of the eukaryotic SENP/ULP family, this interface is mainly stabilized by a large number of backbone hydrogen bonds between the C-terminal tail and NopD (Figure 3a & 3b). The geometry and distances of these contacts between the C-terminal tails of AtSUMO2 and ubiquitin are relatively similar, and almost complete all possible backbone bonds. Singularly, the guanidinium sidechain and carbonyl mainchain of Arg913, located in the “Loop insert” of NopD, contribute with two backbone bonds in ubiquitin and AtSUMO2. Trp836 in NopD sandwiches the C-terminal diGly motif, as usually observed in other members of the SENP/ULP family.

**Figure 3.**
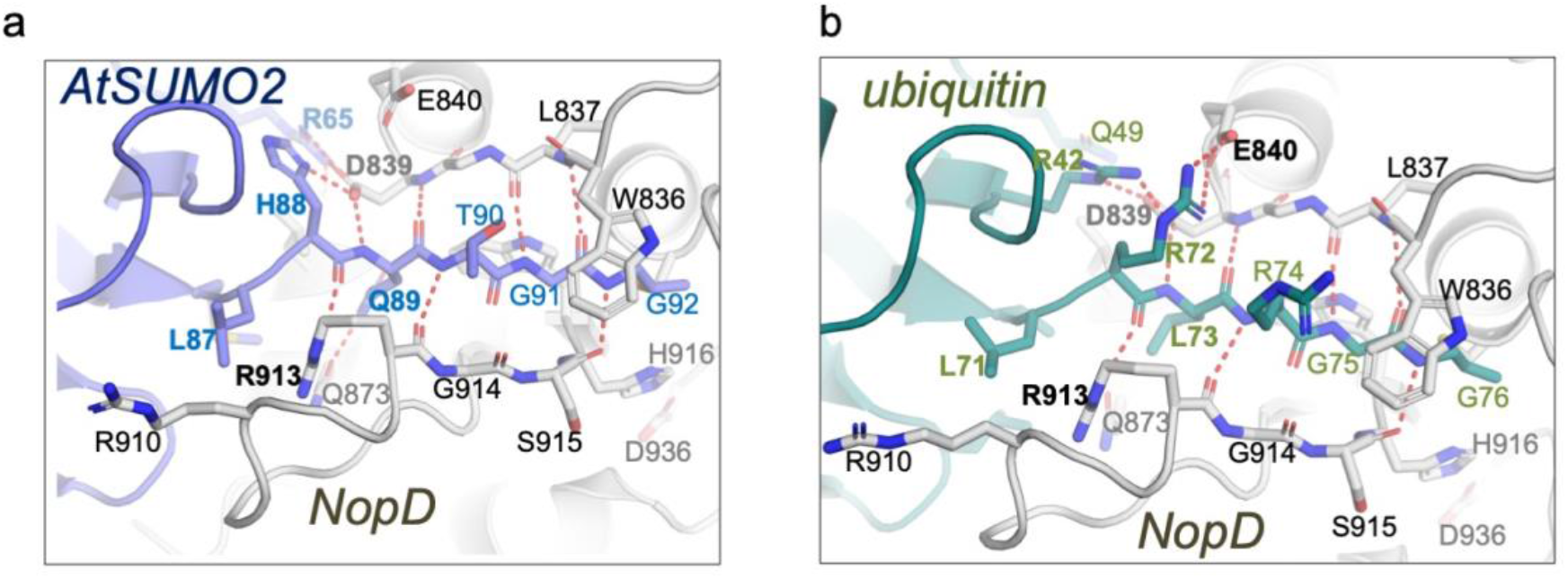
Structural details of the of NopD C-terminal interaction. **a,** Close-up view of the C-terminal tail of AtSUMO2 (purple) in complex with NopD (gray). **b.** Close-up view of the C-terminal of ubiquitin (green) in complex with NopD (gray). Binding side-chain residues in the contact area of substrate and enzyme are shown in stick representation and labelled. Hydrogen bonds are represented by dashed red lines.

All AtSUMO2 and ubiquitin C-terminal side chains are engaged with NopD, except Thr90 or Arg74 in either AtSUMO2 or ubiquitin, respectively (Figures 3a & 3b). Glu89 in AtSUMO2 forms a hydrogen bond with Gln873, which is not present in the equivalent Leu73 in ubiquitin. In vitro assays with NopD Q873N point mutant confirms the role of this hydrogen bond in AtSUMO2, but not in ubiquitin (Figure 5). Interestingly, whilst His88 in AtSUMO2 forms an electrostatic bond with Asp839, the equivalent Arg72 in ubiquitin binds Glu840 (Figures 3a & 3b). Moreover, Leu87 and Leu71, in AtSUMO2 and ubiquitin, respectively, are located in similar hydrophobic pocket of the “Loop insert” of NopD (Figures 3a & 3b).

A previous study reported that NopD was specific for plant SUMO isoforms ^22^, such as AtSUMO1 and AtSUMO2, GmSUMO (soy bean), PvSUMO (common bean), all containing a similar C-terminal sequence (-MLHQTGG), in contrast to the lack of activity for human SUMO isoforms, which contain different C-terminal sequences (-YQEQTGG in hSUMO1 or -FQQQTGG in hSUMO2/3). The structure of the AtSUMO2-NopD complex sheds light into the SUMO ortholog specificity in plants by the unique contacts established by Leu87 and His88 with NopD residues (Figure 3a), in particular by the electrostatic interaction of AtSUMO2 His88 with NopD Asp839, which cannot be established by the equivalent Glu or Gln in human SUMO1 and SUMO2/3 counterparts, respectively (Figure 1d).

### SUMO2 and ubiquitin interface with NopD

As initially reported in the yeast ULP1-SUMO structure ^34^, an extended interface between the globular domain of SUMO and ubiquitin with NopD is observed. To analyze their specific contacts, the interfaces have been divided in three orthogonal views (Figure 4).

**Figure 4.**
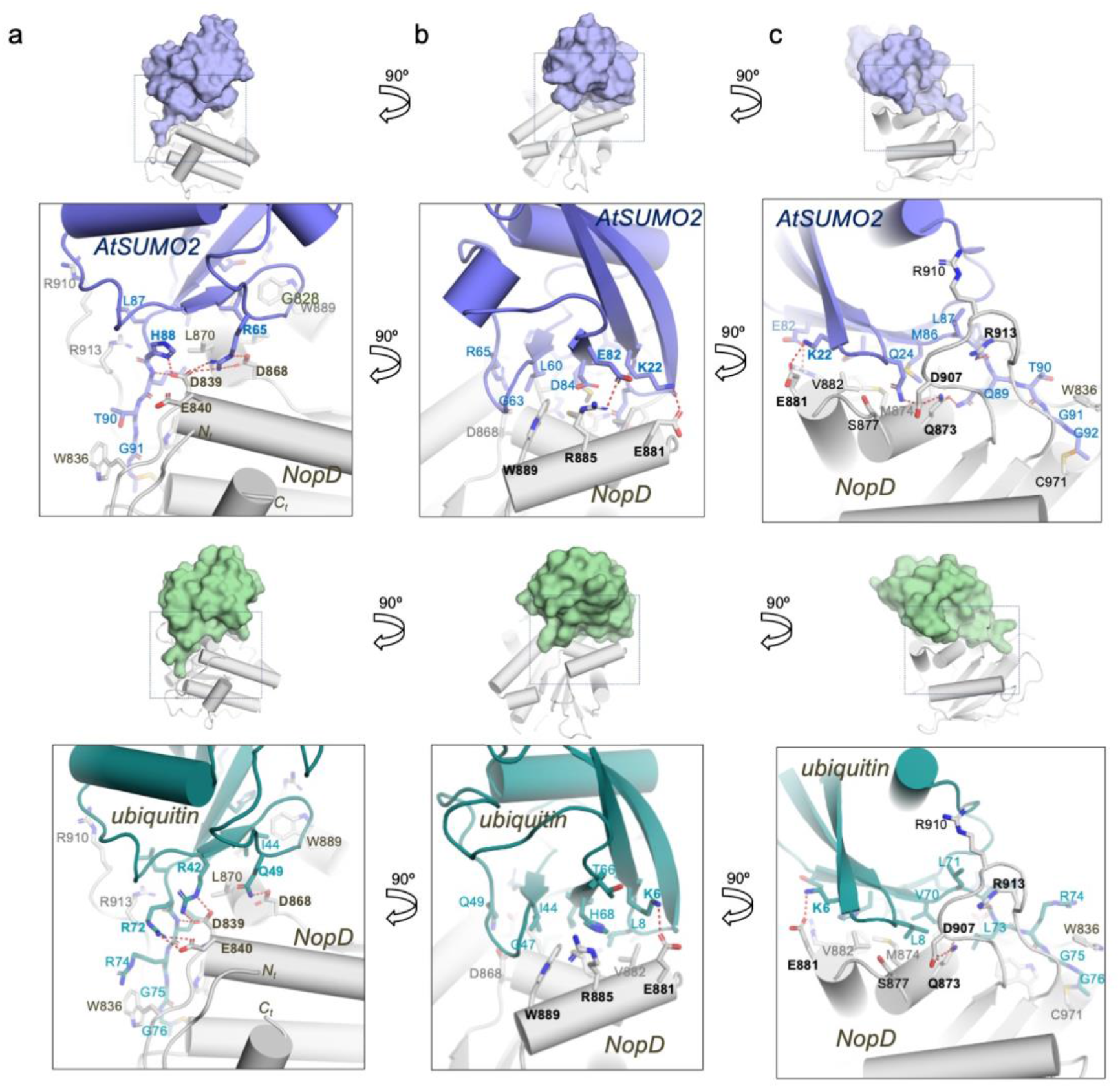
Interface between ubiquitin and AtSUMO2 globular domain with NopD. **a-c,** Close-up view of the ribbon representation of NopD-AtSUMO2 (above) and NopD-ubiquitin (below) interfaces in three orthogonal views. Binding side-chain residues in the contact area are labelled and shown in stick representation. AtSUMO2 and ubiquitin residues are colored in blue and green, respectively. Hydrogen bonds and charged interactions are represented by dashed red lines.

In the first view AtSUMO2 and ubiquitin are engaged to an acidic NopD surface composed by Asp839, Glu840 and Asp868 (Figure 4a). In the AtSUMO2 complex, Asp839 and Asp868 form an electrostatic interactions with His88 and Arg65, respectively, whereas in ubiquitin they interact with Arg42 and Gln49 (in addition to the aforementioned electrostatic interaction between Glu840 and the ubiquitin’s Arg72). It is worth mentioning that only NopD Asp839 is conserved in the eukaryotic SENP/ULP family (Supplementary Figure 1) and, it is essential in the in vitro assays of both SUMO and ubiquitin (Figure 5). However, Glu840 and Asp868 display opposite results in assays with SUMO and ubiquitin. The electrostatic interaction of Glu840 with Arg72 is important in ubiquitin, but not in SUMO, whereas the Asp868 electrostatic interaction with Arg65 is essential in SUMO, but not for ubiquitin (Figure 5).

**Figure 5.**
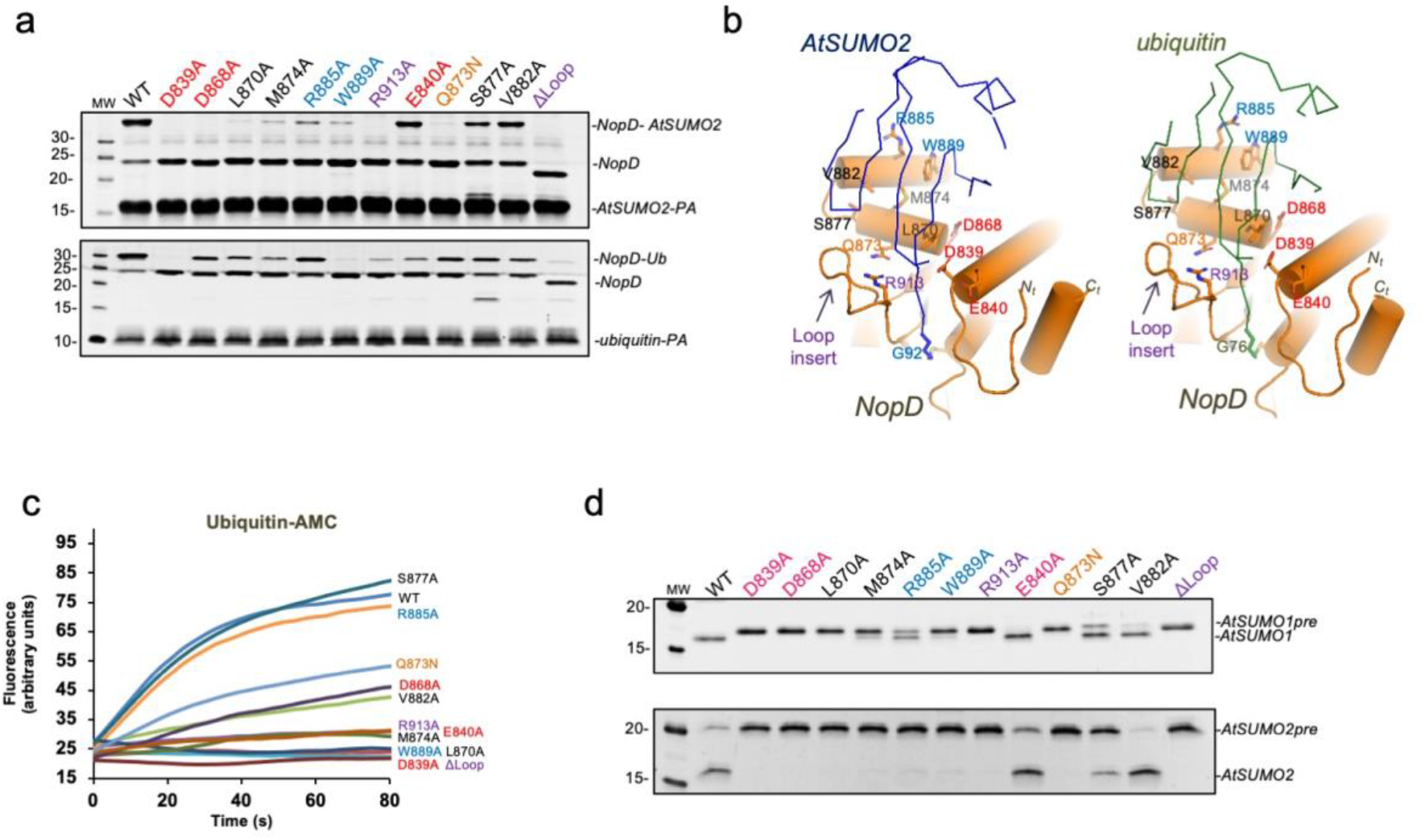
In vitro catalytic analysis of the NopD SUMO/ubiquitin specificity. **a,** SDS-PAGE binding analysis of NopD wild-type and point mutants with AtSUMO2-PA (above) and ubiquitin-PA (below) propargylamine-derived probes. Reaction assays were performed with NopD at 1 μM using the PA probes at 4 μM for 2 hours. **b,** Cartoon representation of the NopD-AtSUMO2 and ubiquitin complexes. AtSUMO2 (blue) and ubiquitin (green) are shown in ribbon representation and NopD (orange) in cartoon representation. Analyzed interface residues are labeled and shown in stick representation. **c,** Time-course plot of the hydrolysis of the fluorescent ubiquitin-AMC substrate with NopD wild type and point mutants. Reactions were conducted in triplicate and the average curve is displayed. **d,** SDS-PAGE of end point activity assays for NopD (200 nM) using the substrates precursor of *Arabidopsis* SUMO1 and SUMO2 (1 μM).

In the second interface view, orthogonal to the C-terminus (Figure 4b), the major contacts emerge from helix α4 are Trp889, Arg885 and Glu881. In ubiquitin, Trp889 is located in a hydrophobic pocket formed by Ile44, Gly47 and His68 (all distances around 3.5 Å); whereas in AtSUMO2, Trp889 is located in a similar pocket formed by Leu60, Gly63 and Asp84 (at distances around 3.5 Å). Mutagenesis analysis highlights the essential role of Trp889 in binding and activity assays for both SUMO and ubiquitin (Figure 5), which is conserved and essential in the eukaryotic SENP/ULP family (Supplementary Fig. 1). Arg885 is engaged in an electrostatic interaction with Glu82 in AtSUMO2, whereas in ubiquitin does not interact with the equivalent His68 (Figure 4b). In vitro assays confirms the major role of Arg885 in SUMO, but not in ubiquitin (Figure 5). Finally, Glu881 is at contact distance to the equivalent Lys22 or Lys6 in AtSUMO2 or ubiquitin, respectively, but the electron density maps do not indicate a strong interaction.

In the third interface view, contacts are established by the unique NopD “Loop insert” with residues emerging from helix α3-α4 loop (Figure 4c). In AtSUMO2 Gln24 and Met86 are placed in a NopD pocket formed by Leu870, Gln873 and Met874 from helix α3 and Val882 from helix α4, whereas in ubiquitin the equivalent Leu8 and Val70 interact with a similar NopD pocket. Asp907, from the NopD “Loop insert”, forms a hydrogen bond with Gln24 in AtSUMO2. Ser877 in the α3-α4 loop forms a strong hydrogen bond with the mainchain of Gln24 or Leu8 in either SUMO or ubiquitin, respectively (around 2.7 Å distance). However, the NopD S877A point mutant does not display a major effect in in vitro assays with SUMO or ubiquitin (Figure 5).

### Dissecting the deconjugation activities in tobacco leaves infiltration assays

The deconjugating activity of NopD was shown to be essential in the induction of cell death after *Agrobacterium* infiltration in leaves of tobacco plants ^22^. We have recapitulated these experiments using full-length and catalytic domain constructs of NopD wild-type, C971A and E840A single point mutants. Our aim was to distinguish between the role of the different deconjugation activities in NopD, SUMO and ubiquitin, using a single point mutant that affects only the deubiquitinating activity in NopD, namely E840A (Figure 5).

Infiltration experiments with the NopD catalytic domains display an equivalent cell death and photosynthetic efficiency between C971A and E840A point mutants in *Agrobacterium* infiltrations of tobacco leaves (Figure 6a). These results indicate that the cell death phenotype in tobacco plant leaves might be basically caused by a loss of the NopD deubiquitinating activity, displaying comparable results between global deconjugation loss (C971A mutant) and loss of deubiquitinating activity (E840A mutant). However, in tobacco leaves infiltrations with the NopD full length, including the long unstructured N-terminal extension, cell death and photosynthetic efficiency are more evident after a shorter period, and in this instance the NopD E840A point mutant shows a stronger phenotype than in the catalytic domain alone, but still lower compared to the NopD wild-type infiltrations (Figure 6b).

**Figure 6.**
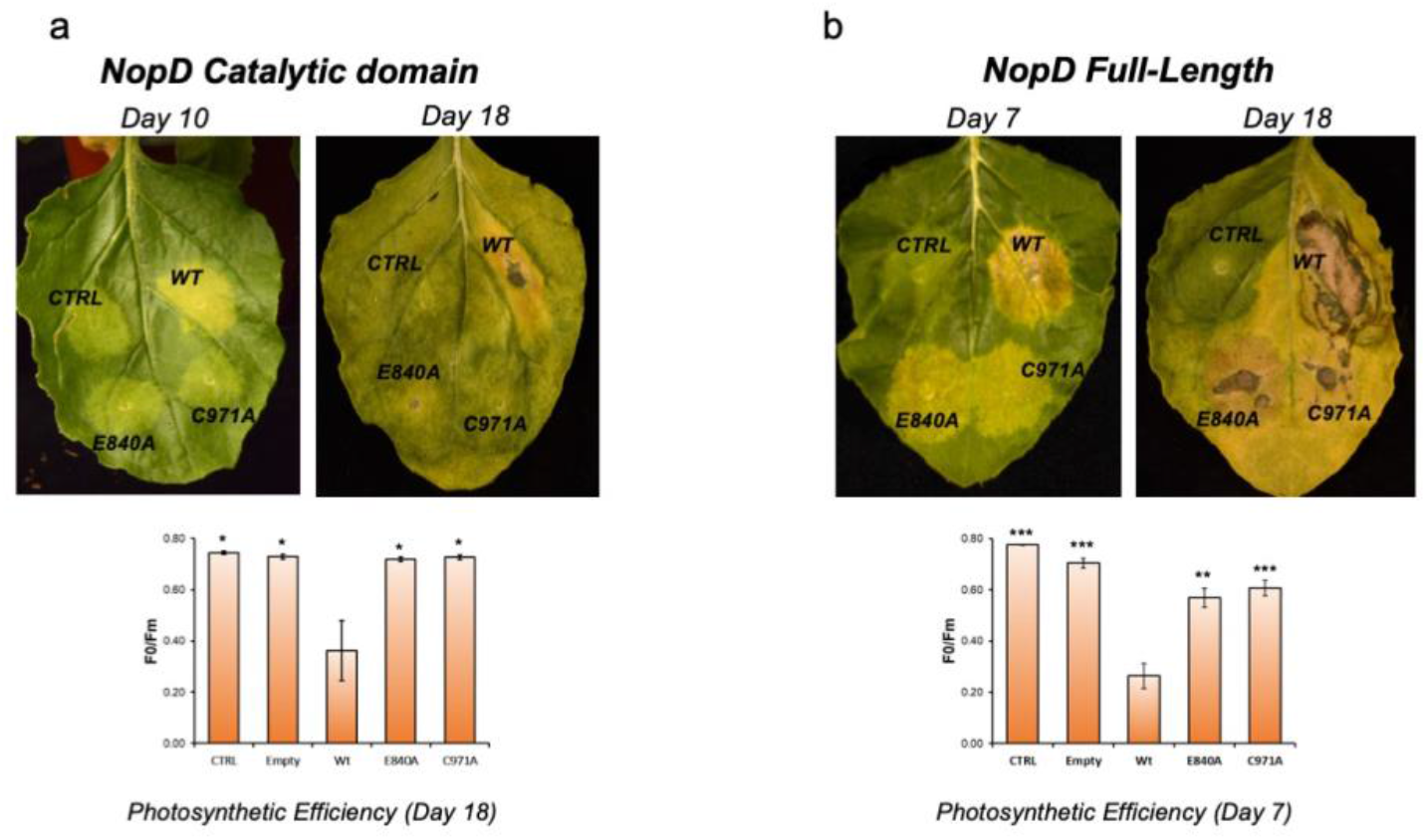
Expression of NopD and NopD mutants in plant cells induces cell death. Expression of **a,** catalytic domain or **b,** full-length NopD WT, NopD C971A and NopD E840A proteins under the control of CAMV 35S promoter in different sections of a leaf was performed by infiltration of *A. Tumefaciens* transformed with pCAMBIA plasmids carrying the described NopD genes. Empty plasmid was used as a control for comparison. The photographs were taken during several days post infiltration (dpi). Cell death (necrotic tissue, can be visualized in wtNopD (yellow-brown areas), whereas expression of NopD C971A or NopD E840A showed no visible effects (**a,b top**). Quantification of cell death was measured by the reduction in photosynthetic efficiency of the tissue surrounding the injection point on each section of the leaf (**a,b bottom**). Measurements around the injection point were taken for each section using five independent leaves. Therefore, data values represent the mean ±SD, n = 5 biological replicates. Significance was measured by a two-tailed unpaired t-test relative to wild type. All data were analyzed with a 95% confidence interval. *P < 0.02, **P < 0.002, ***P < 0.0004. Exact P values from the left to right: (a) 0.011, 0.014, 0.016, 0.014; (b) < 0.0001, < 0.0001, 0.0011, 0.0003.

These results indicate that the long N-terminal unstructured extension of NopD might play a determinant role in the hypersensitive response during infiltration of tobacco leaves, and that in both NopD constructs, catalytic domain and full-length, results point to a stronger dependence on the deubiquitinating activity of NopD for the cell death infection phenotype observed in tobacco leaves.

## Discussion

The eukaryotic CE protease clan in humans, the SENP/ULP family, consists of six SUMO-specific and one Nedd8-specific members. In humans, the conserved catalytic domain has evolved to distinguish between SUMO isoforms and in the case of SENP8/DEN, it has even been adapted to interact with a different UbL modifier, Nedd8 (but not ubiquitin). In infectious bacteria, secreted effector proteins also contain CE proteases to perturb the host response processes. However in bacteria, the CE protease clan has evolved to acquire deubiquitinating activity ^17^, or even an unusual acetyltransferase activities, such as in the *Yersinia pestis* YopJ ^35^. Thus, it seems that in different scenarios the conserved catalytic fold of the CE protease family has evolved to cleave off different types of UbL modifiers.

The particular ability in the CE proteases of pathogenic bacteria to cleave off host-cell ubiquitin conjugates can be accomplished by sequence insertions between conserved secondary structure elements of the catalytic domain ^17^. In eukaryotic SENP/ULP members, sequence insertions are also present to distinguish SUMO isoforms, such as in SENP6 and SENP7 ^36^, or even in NEDP1/SENP8 to cleave Nedd8 ^32,33^. So far all infectious human pathogens show a preference for targeting K63-linkage ubiquitin chains, which are involved in inflammatory signaling cascades. In contrast, the plant pathogen XopD, from *Xanthomonas campestris,* prefers the K48-linkage ubiquitin chains over K63- and K6-linkages, also highlighting the versatility of the CE clan to modulate cleavage of different types of ubiquitin chains. *Xanthomonas* XopD has an unusual dual deconjugation activity for both SUMO and ubiquitin by the formation of an atypical protein interface with ubiquitin through an N-terminal extension.

Here we have discovered that NopD, an effector protease from *Bradyrhizobium* involved in symbiotic plant nodulation, possesses multiple deconjugation activities, including deSUMOylation (*Arabidopsis* SUMO1 and SUMO2), deubiquitination (preferring cleavage of K48-linkage ubiquitin chains), and deNeddylation (Nedd8). Our structural studies reveal that in contrast to *Xanthomonas* XopD, NopD binding to ubiquitin is engaged by the same interface that binds to SUMO, an unexpected feature considering the substantial surface differences between SUMO and ubiquitin. Since NEDD8 and ubiquitin are the two most closely related among all the UbLs, it would be expected that a similar binding interface with NopD. The contrast in ubiquitin binding by the homologues NopD and XopD represent an excellent example of convergent evolution in distant organisms to cleave off ubiquitin by formation of different protein-protein interfaces. Whereas NopD interacts with ubiquitin by sharing a similar interface with SUMO, *Xanthomonas* XopD relies on a particular N-terminal extension that shapes a novel ubiquitin interface.

A particular feature in NopD that allows such multiple deconjugation activities is the presence of a unique insertion, named “Loop insert”, that tethers the C-terminal tail of both SUMO and ubiquitin. Structural and mutagenesis analysis highlights the role of the NopD “Loop insert”, allowing Arg913, which is conserved in the NopD clan (Supplementary Figure 5), to engage through similar backbone hydrogen bonds the C-terminal tails of SUMO and ubiquitin. Interestingly, despite evident differences in their surface residues, *Arabidopsis* SUMO1 and SUMO2 share with ubiquitin common features in the “Loop insert” interaction. In particular Leu87 and His88 in the C-terminus of *Arabidopsis* SUMO1 and SUMO2 structurally align with Leu71 and Arg72 of ubiquitin. Whereas Leu87 (AtSUMO1) and Leu71 (ubiquitin) bind the hydrophobic sidechain segment of Arg910 at the “Loop insert”, the positive charges of His88 (AtSUMO) and Arg72 (ubiquitin) bind Asp839 and Glu840, respectively (Figure 3 and 4). On the other hand, human SUMO1 and SUMO2, with either Gln or Glu/Gln located in those positions (Figure 1g), are unable to bind neither NopD nor the *Xanthomonas* XopD ^23^.

Another peculiar feature in NopD to facilitate ubiquitin binding is the electrostatic interaction between Glu840 and Arg72 from the C-terminal tail of ubiquitin. Whereas in *Xanthomonas* XopD Arg72 interacts with a glutamic acid from the N-terminal extension, in NopD Arg72 is engaged with Glu840 (glycine in *Xanthomonas* XopD) (Figure 2a). In DUBs, such as USPs or OTU families ^33,37,38^, the ubiquitin C-terminal Arg72 has usually been employed as a major binding point to correctly position the ubiquitin C-terminal tail in the active site protease. Interestingly, our functional analysis indicated that the E840A point mutant does not play a significant role in the deSUMOylating activity or binding to AtSUMO2 (Figure 5), in contrast to its relevance in deubiquitinating activity, binding to ubiquitin and ultimately, in the cell death infection phenotype in tobacco leaves (Figure 6). Our results underlies Glu840 as an evolutive trait in *Bradyrhizobium* to enhance binding and cleavage of ubiquitin by NopD.

The cross-reactivity of NopD for SUMO and ubiquitin by using a similar interface is an unusual feature in the CE protease clan. Only in some other human pathogen effectors, such as ChlaDUB1 and RickCE, cross-reactivity can be found for ubiquitin and Nedd8 using a similar interface ^17^, but in this case both UbL modifiers share a 58% sequence identity and an almost identical C-terminal tail (Figure 1d). This multiple deconjugation activity of NopD for SUMO, ubiquitin and Nedd8 is somehow surprising, basically by the presence of a dissimilar C-terminal tail and non-conserved surface residues of the globular domain of these UbL modifiers (Figure 1d). So, it seems that in *rhizobia* NopD has evolved to possess multiple deconjugation activities, at least for SUMO, ubiquitin and Nedd8, in a single polypeptide chain by the acquisition of a “Loop insert” sequence and by maintaining few “hot spots” in the interface necessary for the proteolytic activity of each type of UbL modifiers.

## Materials and methods

### Plasmids, Cloning and Point Mutation

The plasmid for the catalytic domain of NopD was cloned from pMx-NopD (purchased from Thermo Fisher Scientific). All constructs of pET28a-NopD were amplified by PCR and ligated using the “Restriction Enzyme Free PCR” methodology ^39^. The NopD point mutants constructs were generated by different primers and were created by the QuickChange site-directed mutagenesis kit (Stratagene). All primers are shown in the Supplementary Table1. The plasmid of pCAMBIA-NopD Full-length and catalytic domain were cloned from pET28-NopD Full-length (a gift from Dr. Christian Staehelin). *Arabidopsis thaliana* SUMO1 and SUMO2 precursors were from M. Lois lab. All plasmids for the pTXB1-AtSUMO2G and pTXB1-UbiquitinG expression were cloned using the “Restriction Enzyme Free PCR” methodology ^39^.

### Protein expression and purification

The NopD-CD (catalytic domains), *Arabidopsis thaliana* SUMO2 (AtSUMO2) and human ubiquitin expression constructs were transformed into E. coli Rosetta (DE3) cells (Novagen). Bacterial cultures were grown at 37 °C to OD600=0.7∼0.8, and IPTG was added to a final concentration of 0.5 mM. Cultures were grown an additional 5 h at 30 °C and harvested by centrifugation. Cell were suspended in 20 mM Tris-HCl (pH 8.0), 350 mM NaCl, 10 mM imidazole, 20% sucrose, 1 mM DTT, and 0.1% IGEPAL CA-630, and cells were broken by sonication. After cell debris removal by centrifugation, proteins were purified by nickel affinity chromatography using Ni Sepharose 6 Fast Flow (GE Healthcare) and eluted with 20 mM Tris-HCl (pH 8.0), 350 mM NaCl, 300 mM imidazole, and 1 mM DTT. Proteins were further purified by gel filtration (Superdex 75; Cytiva) equilibrated with 20 mM Tris-HCl (pH 8.0), 250 mM NaCl, and 1 mM DTT. Finally, for NopD-CD, GF fractions containing the target protein were pooled, diluted to 50 mM NaCl and, applied to an anion exchange resin (Resource S; Cytiva), and eluted with a 0-1 M NaCl gradient from 0 to 50% in 20 mM HEPES (pH 8.0) and 1 mM DTT. Proteins were then concentrated using Amicon Ultra ultrafiltration device (Milipore) and snap-frozen in liquid nitrogen prior to storage at −80 °C.

### Preparation of the NopD complexes with AtSUMO2 and ubiquitin

The method to generate AtSUMO2-PA or Ub-PA protein derivatives has been previously described ^40^. NopD-CD protein and AtSUMO2-PA/Ub-PA (1:4 ratio) were incubate for 3h at 30 °C. After that, the buffer was changed to 20mM HEPES 7.42, 50mM NaCl, 1mM DTT by rounds of concentration and dilution with Amicon Ultra-30K ultrafiltration device. Complexes were purified using an anion exchange resin (Resource S; Cytiva) as described above.

### Crystallization and Data Collection

NopD-AtSUMO2-PA and NopD-Ub-PA were concentrated to 12 mg/mL for crystallization screening. Crystallization experiments were performed at 18°C by sitting drop vapor diffusion method. NopD-AtSUMO2-PA crystals grew up in a protein mixture with an equal volume of a condition solution containing 0.1 M Imidazole, pH 7.0 and 50% v/v MPD. NopD-Ub-PA crystals grew up in a protein mixture with an equal volume of a condition solution containing 0.1 M Imidazole pH 8.0 and 10% w/v PEG8000. Crystals were harvested after 1-2 weeks and soaked 5-10 seconds in the crystallization buffer supplemented with 15% ethylene glycol, and then snap-frozen in liquid nitrogen to storage.

Diffraction data were collected at beamline BL13-XALOC at the ALBA synchrotron (Barcelona, Spain). NopD-AtSUMO2PA and NopD-UbPA get a resolution: 1.50 Å and 1.94 Å, respectively. Resolution Data processing was conducted by AutoProcesing with MxCUBE ^41,42^. Structures were solved by molecular replacement with NopD AlphaFold2 model as a search mode ^43^. Following rounds of model building and refinement were carried out with Coot and Phenix ^44,45^.

### In vitro de-SUMOyation assays

Protease activity was measured by incubating precursor of *Arabidopsis thaliana* SUMO1/2 with purified 200 nM of NopD wild type and mutants at 30 °C in a buffer containing 20 mM Tris-HCl (pH 8.0), 250 mM NaCl, and 2 mM DTT. SDS-BME loading buffer was used to terminate the reactions after 2 hours, and gel electrophoresis was used to examine the results (PAGE). SYPRO staining was used to identify proteins (Bio-Rad). A Gel-Doc machine and integration software were used to detect and quantify the products (ImageLab; Bio-Rad).

### Activity-based binding probe assays

NopD wild-type and active site mutant (C971A) were prediluted to 1 μM in reaction buffer 20 mM Tris-HCl (pH 8.0), 250 mM NaCl, and 4 mM DTT, and combined 1:4 with 4 μM of Ub-PA, AtSUMO1-PA, AtSUMO2-PA and hSUMO2-PA for 2 hours at 30 °C. The reaction was stopped by adding SDS-PAGE loading buffer and checked by SDS-PAGE.

### AMC-substrate hydrolysis assays

NopD wild type and point mutants were incubated with Nedd8-, ubiquitin-, SUMO1- or SUMO2-AMC at 30°C and measured the fluorescence emission using 345 nm excitation and 445 nm emission wavelengths using a Jasco FP-8200 spectrofluorometer. All measurements were carried out in triplicate with 25 nM NopD and 0.1 μM UbL-AMC substrate in a buffer containing 100mM NaCl, 20 mM Tris-HCl pH 8, 10 mM DTT.

### Cell-Death Tobacco Leaves assays

*Agrobacterium tumefaciens* GV3101 cells were transformed with the constructs pCAMBIA-NopD-CD (Catalytic Domain) or pCAMBIA-NopD-FL (Full length) and grown in YEB plates supplemented with rifampicin (100 μg/ml), gentamycin (50 μg/ml) and Kanamycin (25 μg/ml). Single transformed colonies (Checked by colony PCR) were grown o/n in 3 ml YEB medium supplemented with the same antibiotics at 28°C. Fresh medium (20 mL) was inoculated with the precultures and grown o/n at 28°C. The cultures were centrifuged and collected cells were resuspended in induction buffer (10 mM MgCl_2_, 10 mM MES, 150 μM acetosyringone, pH=5.6) to achieve OD_600_=4. The mixture was incubated at room temperature for 3h and then mixed with the same volume of a culture harboring vector pGWB702-HCProWMV at the same OD to prevent silencing ^46^ and induction buffer for a final OD=0.5 for each strain. Mixtures were infiltrated in two differents leaves of 5-10 3-4-week-old *N. benthamiana* plants grown under normal greenhouse conditions as described ^46^. Agroinfiltrated leaves were collected at sequential days post-infiltration (dpi) and phenotypes recorded using a Camera NIKON D7000. Cell-death was quantified as the loss of photosynthetic capacity in the infiltrated regions of the leaves by measuring the *In vivo* Chl fluorescence at room temperature using a pulse modulated amplitude fluorimeter (MAXI-IMAGING-PAM, Heinz Waltz GmbH, Germany). Photosynthetic parameter of variable to maximum fluorescence (*F*v/*F*m) weas measured as described elsewhere ^47^. Briefly, Leaves were first dark-adapted for 30 min and then a very short saturating pulse (SP) was applied. Fluorescence signal before (*F*o) and after (*F*m) the SP was recorded to estimate the maximum quantum yield of photosystem II (PSII) (*F*v/*F*m).

## Acknowledgements

This work was supported by grants from the *“Ministerio de Ciencia e Innovación*” PID2021-124602OB-I00, and ICREA, ICREA-Academia-2022 to DR. YL acknowledges her scholarship of the China Scholarship Council program from the Chinese government. DR acknowledges support from the Serra Hunter program from Generalitat de Catalunya. Thanks to Dr. Christian Staehelin from Sun Yat-sen University for NopD Full-length plasmid. X-ray experiments were performed at BL-13 XALOC beamline at ALBA Synchrotron with the collaboration of ALBA staff.

## Author contributions

YL conducted the crystallization experiments. YL and NV conducted in vitro activity assays. JPG and ML conducted the plant infiltrations. NV and DR contributed to the correction and writing of the manuscript.

