## Supplementary Figures for "Multipotent ubiquitin/ubiquitin-like deconjugation activity of the *rhizobial* effector NopD"

```
                                                              α1                α2
      790                                     828     840
NopD  LGPRGRRVAQFIHHPRPSPVPGAQIGPSATVASSGHRSGPVLGPTQWLGDEHIQRDYELLAQELQQNNPDLAAR
XopD  -EYPNNHGTQTPMGYSAMTPERIDVDNLPSPQDVADPELPPVRATSWLLDGHLRAYTDDLARRLRG-EPNAHL-
SENP1 EEMEKEIKNVFRNGNQDEVLSEA------FRLTITRKDIQTLNHLNWLNDEIINFYMNMLMERSK--EKGLPS-
SENP2 EDMEKEISNALGHGPQDEILSSA------FKLRITRGDIQTLKNYHWLNDEVINFYMNLLVERNK--KQGYPA-
SENP3 LQLIQSYQRMPGNAMVRGFRVAY------KRHVLTMDDLGTLYGQNWLNDQVMNMYGDLVMDTVP--EK-----
SENP5 PFINREITNYRARHQKCNFRIFY------NKHMLDMDDLATLDGQNWLNDQVINMYGELIMDAVP--DK-----
SENP6 DEEEETGENHTIFIGPVEKLIVYPPPPAKGGISVTNEDLHCLNEGEFLNDVIIDFYLKYLVLEKL--KKEDADR
SENP7 TSNPDEEWREVRHTGLVQKLIVYPPPPTKGGLGVTNEDLECLEEGEFLNDVIIDFYLKYLILEKASDELVER--
SENP8 ---------------MDPVVLSY------MDSLLRQSDVSLLDPPSWLNDHIIGFAFEYFANS-QFHDCSDH--

       β1    α3                     α4                 β2       Loop Insert   β3
      864                                 889                              916
NopD  TRFVDPLIAQMLRS-------------PSKEVAERALGWVRP---GTAD-FLFLPVSDASDTDRHQRGSHWSLL
XopD  LHFADSQVVTMLSS-------------ADPDQQARAQRLLAG---DDIPPIVFLPINQPN--------AHWSLL
SENP1 VHAFNTFFFTKLKTA-----------------GYQAVKRWTKKVDVFSVD-ILLVPIHLGV---------HWCLA
SENP2 LHVFSTFFYPKLKSG-----------------GYQAVKRWTKGVNLFEQE-IILVPIHRKV---------HWSLV
SENP3 VHFFNSFFYDKLRTK-----------------GYDGVKRWTKNVDIFNKE-LLLIPIHLEV---------HWSLI
SENP5 VHFFNSFFHRQLVTKG-----------------YNGVKRWTKKVDLFKKS-LLLIPIHLEV---------HWSLI
SENP6 IHIFSSFFYKRLNQRER-RNHETTNLSIQQKRHGRVKTWTRHVDIFEKD-FIFVPLNEAA---------HWFLA
SENP7 SHIFSSFFYKCLTRKENNLTEDNPNLSMAQRRHKRVRTWTRHINIFNKD-YIFVPVNESS---------HWYLA
SENP8 VSFISPEVTQFIKCTSN--------------PAEIAMFLEPLDL-PNKR-VVFLAINDNSNQAA--GGTHWSLL

       β3                    β4              α5                       β5
      920                      936
NopD  LVDRRDR------------GRRVAYHYDSTQ-----GYNDGLAAELAGRLD------------ANLQQAP----
XopD  LVDRRNK------------DAVAAYHYDSMA------QKDPQQRYLADMAAYHLGLDY-----QQTHEMP----
SENP1 VVDF---------------RKKNITYYDSMGGIN-NEACRILLQYLKQESIDKKRKEFDT--NGWQLFSKKSQE
SENP2 VIDL---------------RKKCLKYLDSMGQKG-HRICEILLQYLQDESKTKRNSDLNLL--EWTHHSMKPHE
SENP3 SVDV---------------RRRTITYFDSQRTLN-RRCPKHIAKYLQAEAVKKDRLDF---HQGWKGYFKMN--
SENP5 TVTL---------------SNRIISFYDSQGIHF-KFCVENIRKYLLTEAREKNRPEF---LQGWQTAVTKC--
SENP6 VVCFPGLEKPKYE//PTICKQPCILLMDSLRGPSRSNVVKILREYLEVEWEVKKGSKRSFSKDVMKGSNPK---
SENP7 VICFPWLEEAVYE//KKMCKRPCILILDSLKAASVQNTVQNLREYLEVEWEVKLKTHRQFSKTNMVDLCPK---
SENP8 VYLQ---------------DKNSFFHYDSHSRSNSVHAKQVA-EKLEAFLGRKGDKLAF------VEEK-----

                     α6                       α7
      962      971                                          1016
NopD  IRQQQNSYDCGVFVLDGTRELVRRLAARRPDL----NLNNLVISRQELRDRLGAGVGFN
XopD  IAIQSDGYSCGDHVLTGIEVLAHRVLDGTFDYAGGRDLTDIEPDRGLIRDRLAQAEQAPAESSIRQVP
SENP1 IPQQMNGSDCGMFACKYADCITKDRPI-----------NFTQQHMPYFRKRMVWEILHRKLL
SENP2 IPQQLNGSDCGMFTCKYADYISRDKPI-----------TFTQHQMPLFRKKMVWEILHQQLL
SENP3 VARQNNDSDCGAFVLQYCKHLALSQPF-----------SFTQQDMPKLRRQIYKELCHCKLTV
SENP5 IPQQKNDSDCGVFVLQYCKCLALEQPF-----------QFSQEDMPRVRKRIYKELCECRLMD
SENP6 VPQQNNFSDCGVYVLQYVESFFENPILSFELPMNLAN-WFPPPRMRTKREEIRNIILKLQEDQSKEKRK
SENP7 VPKQDNSSDCGVYLLQYVESFFKDPIVNFELPIHLEK-WFPRHVIKTKREDIRELILKLHLQQQKGSSS
SENP8 APAQQNSYDCGMYVICNTEALCQNFFRQQTE-------SLLQLLTPAYITKKRGEWKDLITTLAKK
```

**Supplementary Figure 1. Structural alignment of the *Bradyrhizobium* NopD catalytic domain with the catalytic domains of *Xanthomonas campestris* XopD, and the human SENP/ULP family members: SENP1, SENP2, SENP3, SENP5, SENP6, SENP7 and SENP8/NEDP1.** A yellow backdrop indicates side chain homology with 80% conservation in the alignment. Red represents the three active site catalytic residues. NopD secondary structures (α-helix and β-strands) are numbered and labelled.

a b

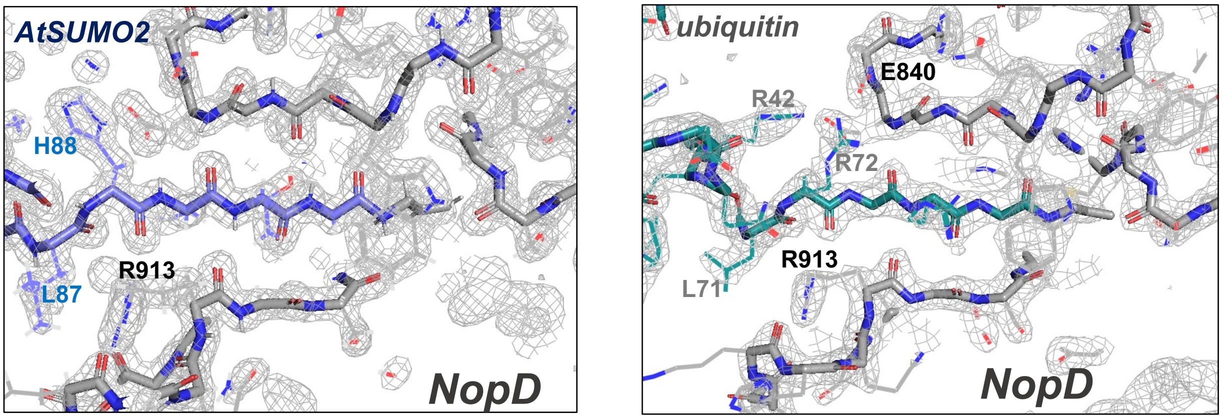


**Supplementary Figure 2. Electron density maps of the C-terminal tails.** Detail of the 2FoFc electron density maps around the C-terminal region of AtSUMO2/Ubiquitin in complex with NopD (grey), contoured at 2σ. Backbone chain is shown in stick representation, whereas side chains are represented in line. Some binding residues are labelled.

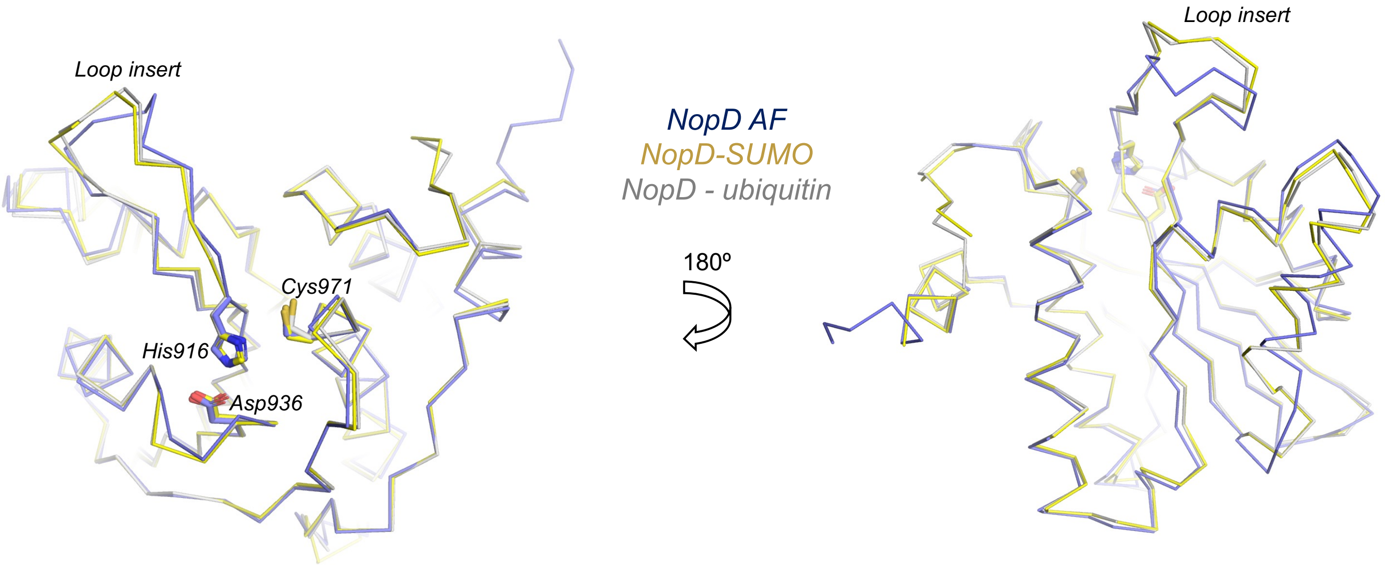


**Supplementary Figure 3. Structural alignment of the catalytic triad residues of NopD in the complex structures.** Two views of the overlapping ribbon structure of the unbound NopD AlphaFold model (blue), NopD-AtSUMO2 complex (yellow) and NopD-ubiquitin complex (grey). SUMO2 and ubiquitin has been removed from the picture. Location of the "Loop insert" is marked. Catalytic residues are labeled and shown in stick representation.

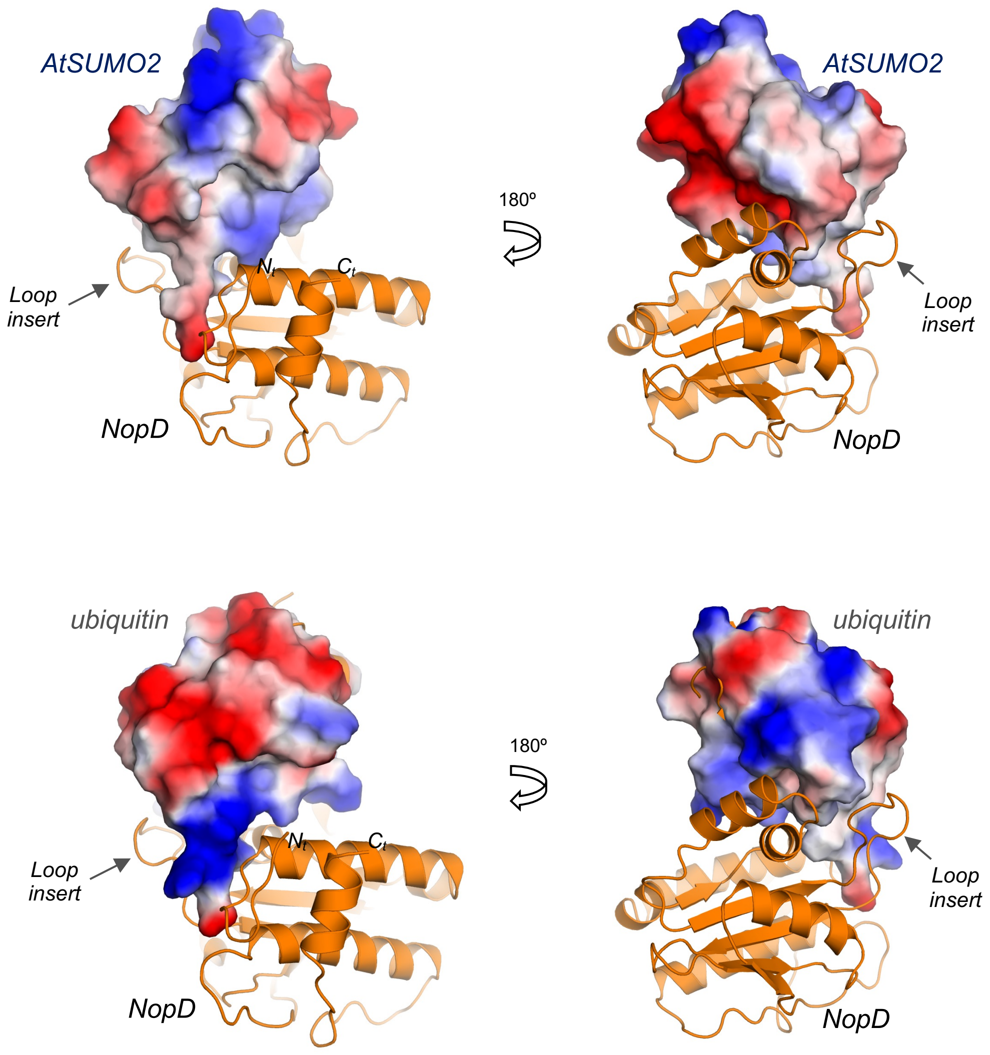


**Supplementary Figure 4. Electrostatic potential surface representation of AtSUMO2 and ubiquitin in complex with NopD.** Two views of electrostatic potential surface representation for AtSUMO2 and ubiquitin in complex with NopD. Red and blue colours indicate negative and positive charges, respectively. NopD is shown in an orange cartoon representation. N-terminal end, C-terminal end and Loop insert are marked and labelled.

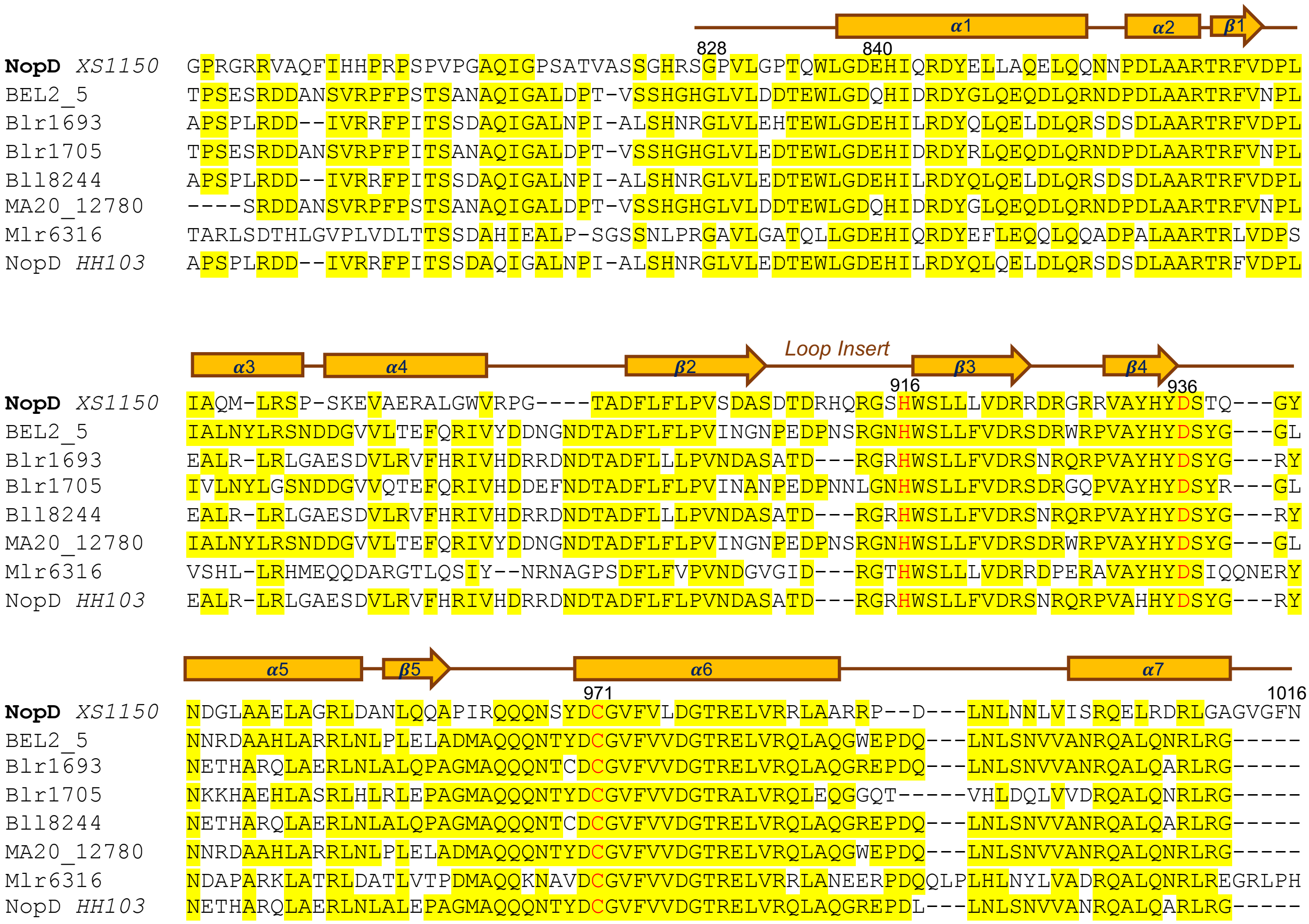


**Supplementary Figure 5. Sequence alignment of the conserved catalytic domains of NopD-related proteins**. Catalytic triad residues are marked in red. A yellow backdrop indicates 80% identity in the alignment. NopD secondary structures (α-helix and β-strands) are numbered and labelled in orange. NopD XS1150 is from *Bradyrhizobium* sp. XS1150, GenBank: MF100854.1; BEL2_5 is form *Bradyrhizobium elkanii* USDA 61, GenBank: AKS25901.1; Blr1693 is from *Bradyrhizobium diazoefficiens* USDA 110, GenBank: BAC46958.1; Blr1705 is from *Bradyrhizobium diazoefficiens* USDA 110, GenBank: BAC46970.1; Bll8244 is from *Bradyrhizobium diazo efficiens* USDA 110, GenBank: BAC53509.1; MA20_12780 is from *Bradyrhizobium japonicum*, GenBank: KGT79298.1; Mlr6316 is from *Mesorhizobium japonicum* MAFF 303099, GenBank: BAB52630.1; NopD HH103 is from *Sinorhizobium fredii* HH103, GenBank: CCE98838.1.

| | |
|---|---|
| NopD 833 fw | CTGGTGCCGCGCGGCAGCCATGCGACACAGTGGCTGGGTGATGAACATATT |
| NopD 833 rv | GTCGACGGAGCTCGAATTCGGATCCTTAGTTAAAACCAACACCGGCACCT |
| NopD 827 fw | CAGGATACAGTCTAACCAAGCGAGAGGATAAGCATTTTTCC |
| NopD 791 fw | TGGTGCCGCGCGGCAGCCATGGTCCTCGTGGTCGTCGTGTT |
| NopD 798 fw | CTGGTGCCGCGCGGCAGCCATGCACAGTTTATTCATCATC |
| PTXB1-AtSUMO2Gfw | ACTTTAAGAAGGAGATATACATATGTCTGCTACTCCGGAAGAAGACAAG |
| PTXB1-AtSUMO2G rv | AGTGCATCTCCCGTGATGCAACCAGTCTGATGAAGCATTGCAT |
| ∆ A108-G117 Fw kpnI | GATTTTCTGTTTCTGCCGGTTAGTGATGGTACC |
| ∆ A108-G117 Rv kpnI | GATCAACCAGCAGCAGGCTCCAATGGCTGGTACC |
| NopD D42A fw | GTCCGACACAGTGGCTGGGTGCTGAACATATTCAGCGTGA |
| NopD D42A rv | TCACGCTGAATATGTTCAGCACCCAGCCACTGTGTCGGAC |
| NopD D71A fw | GCACGTACCCGTTTTGTTGCTCCGCTGATTGCACAGATGCTGCGTA |
| NopD D71A rv | TACGCAGCATCTGTGCAATCAGCGGAGCAACAAAACGGGTACGTGC |
| NopD Leu73A fw | GTACCCGTTTTGTTGATCCGGCGATTGCACAGATGCTGCGTA |
| NopD Leu73A rv | TACGCAGCATCTGTGCAATCGCCGGATCAACAAAACGGGTAC |
| NopD M77A fw | GTTGATCCGCTGATTGCACAGGCGCTGCGTAGCCCGAGCAA |
| NopD M77A rv | TTGCTCGGGCTACGCAGCGCCTGTGCAATCAGCGGATCAAC |
| NopD R88A fw | GAGCAAAGAAGTTGCAGAAGCCGCATTAGGTTGGGTTCGTCCGGGTA |
| NopD R88A rv | TACCCGGACGAACCCAACCTAATGCGGCTTCTGCAACTTCTTTGCTC |
| NopD W92A fw | GTTGCAGAACGCGCATTAGGTGCGGTTCGTCCGGGTACAGCAGA |
| NopD W92A rv | TCTGCTGTACCCGGACGAACCGCACCTAATGCGCGTTCTGCAAC |
| NopD R116A fw | GCGATACCGATCGTCATCAGGCTGGTAGCCATTGGAGCCT |
| NopD R116A rv | AGGCTCCAATGGCTACCAGCCTGATGACGATCGGTATCGC |

**Supplementary Table 1**. Primers used in this work.
